# A Vascularized Tumoroid Model for Human Glioblastoma Angiogenesis

**DOI:** 10.1101/2021.02.05.429930

**Authors:** Agavi Stavropoulou Tatla, Alexander W Justin, Colin Watts, Athina E Markaki

## Abstract

Glioblastoma (GBM) angiogenesis is critical for tumor growth and recurrence, making it a compelling therapeutic target. Here, a disease-relevant, vascularized tumoroid *in vitro* model with stem-like features and stromal surrounds is reported. The model is used to recapitulate how individual components of the GBM’s complex brain microenvironment such as hypoxia, vasculature-related stromal cells and growth factors support GBM angiogenesis. It is scalable, tractable, cost-effective and can be used with biologically-derived or biomimetic matrices. Patient-derived primary GBM cells are found to closely participate in blood vessel formation in contrast to a GBM cell line containing differentiated cells. Exogenous growth factors amplify this effect under normoxia but not at hypoxia suggesting that a significant amount of growth factors is already being produced under hypoxic conditions. Under hypoxia, primary GBM cells and umbilical vein endothelial cells are found to strongly co-localize in a mosaic pattern to form sprouting vascular networks, with GBM cells acquiring an endothelial-like behaviour, which has been reported to occur *in vivo*. These findings demonstrate that our 3D tumoroid *in vitro* model exhibits biomimetic attributes that may permit its use as a preclinical model in studying microenvironment cues of tumor angiogenesis.

## 1. Introduction

Glioblastoma (GBM) is the most common and aggressive form of brain cancer [1], characterized by significant cell heterogeneity [2], self-renewing cancer stem cells [3], and rich microvasculature [4]. Tumor progression is intimately linked with changes to the tumor microenvironment, including modulation of the extracellular matrix (ECM) and basement membrane compositions [5, 6], disruption of the blood brain barrier [7], changes to matrix moduli and density [8, 9], and inducing a wide range of interactions between endothelial and stromal cell populations [10, 4]. One of most significant features of GBM is its hypervascularity and there is a significant correlation between the degree of angiogenesis and prognosis [11, 12]. A significant challenge exists to accurately recreate this neoangiogenesis and the tumor microenvironment through personalised *in vitro* disease models, which would support the development of new anti-angiogenic treatments to GBM.

Hypoxic and necrotic conditions characterize the GBM tumor microenvironment [13] and lead to complex mechanisms supporting the formation of new vessels and recruitment of immunosuppressive cells [14]. Through HIF-1 signalling, these hypoxic and necrotic niches activate and enrich glioblastoma stem cells (GSCs), induce the transdifferentation of GSCs into endothelial cells, and upregulate the release of pro-angiogenic growth factors, such as vascular endothelial growth factors (VEGF) and basic fibroblast growth factor (bFGF) [4]. Notably, tumors that have a high stem cell number are highly angiogenic [15, 16]. Alongside xenograft and genetically engineered transplant models [17, 18], there is a significant need for 3D *in vitro* models which can be studied under highly controllable conditions (e.g. modulating matrix properties, cell population and signalling factor compositions), for basic research and for rapid drug testing. Furthermore, 3D cell culture models outperform 2D models at replicating the *in vivo* tumor processes [19, 20]. To this end, 3D aggregates of tumor cells (i.e. tumoroids) can be generated *in vitro* which exhibit key biomimetic properties including hypoxic responses, interactions with the ECM microenvironment, heterogeneous stroma cells, molecular signal gradients and differential nutrient and waste transport [21]. While several 3D human GBM models have been investigated, most notably looking at the ECM composition, organization, and drug resistance and penetration [21], there is a need for the development of tumoroid models which mimic the dynamics of tumor angiogenesis, given the importance vascularization plays in glioblastoma progression.

Here, we introduce a 3D vascularized tumoroid *in vitro* model using patient-derived primary GBM cells. This model is used to recapitulate how individual components of the GBM’s complex brain microenvironment, such as low oxygen tension (e.g. 1% O_2_), vasculature-related stromal cells (e.g. human umbilical vein endothelial cells (HUVEC), human dermal fibroblasts (HDF) and growth factors (e.g. VEGF, bFGF), support the GBM angiogenesis.

## 2. Results and Discussion

The vascularized tumoroid model is based on spheroids, in which HUVEC and patient-derived primary GBM are in direct contact with each other, supported by HDF embedded in a fibrin gel (Figure 1A). The NCH82 GBM cell line is used for comparison. Variants of the model include GBM-only or HUVEC-only spheroids surrounded by HDF, GBM or no cells in the fibrin gel.

**Figure 1.**
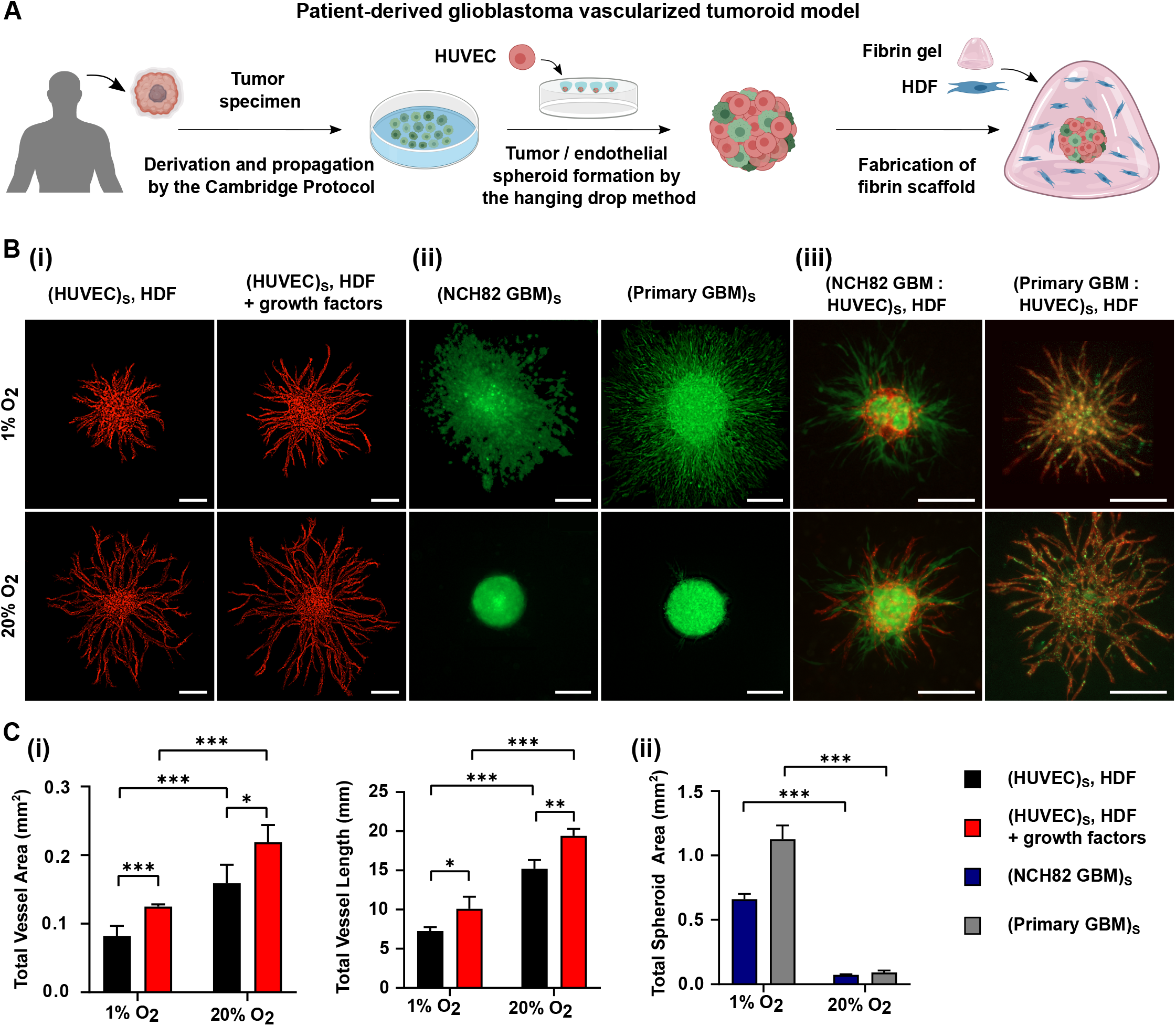
(A) Schematic representation of the formation of a vascularized tumoroid model from patient-derived primary glioblastoma cells. (B) Fluorescence images showing angiogenic sprouting at 1% and 20% O_2_ at day 3 after seeding of (i) HUVEC (red) spheroid, supported by single HDF cells with or without additional growth factors; (ii) NCH82 (green) or primary (green) GBM spheroids and (iii) the full vascularized tumoroid model containing NCH82 or primary GBM: HUVEC spheroids with HDF supporting cells. HDF cells are not stained so they are non-fluorescent. The spheroids and single HDF cells are embedded in a 7.5 mg/mL fibrin gel. The scale bars represent 250 *µ*m. (C) Graphical representations of (i) the total vessel area and vessel length for HUVEC spheroids surrounded by HDF, with or without additional growth factors and (ii) the total spheroid area for NCH82 or primary GBM spheroids. Data are presented as the mean ± standard deviation; *n* = 3; \**p <* 0.05, \*\**p <* 0.01, \*\*\**p <* 0.001. Subscript S denotes cells seeded in the same spheroid.

### 2.1. Sprouting of HUVEC spheroids

The first challenge towards the development of the vascularized tumoroid model is to provide a suitable environment for the HUVEC spheroids to form capillary sprouts. Angiogenesis is a complex process involving dynamic interactions between the endothelial and stromal cells, the production of angiogenic signalling factors, and is affected by changes in the surrounding ECM (e.g. hypoxic conditions). Fibrin was chosen as the supporting matrix for our experiments because it can be derived directly from the patients’ blood [22]. Variation in the ECM composition is a hallmark of tumor stroma, with increased fibrin deposition associated with tumor angiogenesis [23]. Pro-angiogenic factors, such as VEGF and bFGF, have a high affinity for fibrin through the heparin binding domain [24]. Fibrin also triggers the production of other basement membrane proteins that are strongly linked to angiogenesis such as laminin and collagen IV [25].

In the absence of a supporting cell type, endothelial cells cultured in spheroids within a fibrin gel undergo a migratory response instead of forming vasculature (Figure S1). In angiogenesis models, a range of supporting cells can be used including human dermal fibroblasts, mesenchymal stromal cells and pericytes. In this study, we used the well-known method of culturing endothelial cells with human dermal fibroblasts, as they undertake significant collagen deposition and produce pro-angiogenic growth factors enabling capillary formation [26]. To this end, HUVEC spheroids are cultured with HDF single cells (at 10^6^ cells/mL) within a 7.5 mg/mL fibrin gel to form angiogenic sprouts. Experiments are performed under normoxia (20% O_2_) or hypoxia (1% O_2_), with or without additional pro-angiogenic growth factors (200 ng/mL VEGF and bFGF). Figure 1B(i) shows a typical HUVEC spheroid at day 3 after seeding (additional timepoints are shown in Figure S2). In all cases the HUVEC display an angiogenic-like response, sprouting radially outwards forming capillaries. The addition of exogenous growth factors, as well as normoxia, induce a stronger angiogenic response compared to hypoxia. Figure 1C(i) shows the increase in vessel area and total vessel length of the produced sprouts at day 3.

### 2.2. Invasion of GBM spheroids

Before incorporating the tumor cells into the sprouting spheroids, the invasion capability of GBM-only spheroids embedded in a fibrin gel was examined for primary and NCH82 GBM cells under hypoxic and normoxic conditions. Figure 1B(ii) shows fluorescence images of the GBM spheroids at day 3. The corresponding total spheroid areas are shown in Figure 1C(ii). It can be seen that hypoxia significantly increases GBM spheroid area in both primary and NCH82 cells which has been previously reported [27]. This effect is stronger for primary cells compared to NCH82 cells.

### 2.3. Vascularized tumoroid model

The vascularized tumoroid model was created by applying vascularization techniques to tumor spheroids which were presented in the previous section. This model comprises of 1000-cell spheroids, consisting of HUVEC and GBM cells at a ratio of 3:1, in a 7.5 mg/mL fibrin gel containing 10^6^ HDF cells/mL. Both primary and NCH82 GBM cells were used. The primary GBM cells were cultured in serum-free medium that preserves the subpopulation of glioblastoma stem-like cells, whereas the NCH82 GBM cell line was not, resulting in differentiated cells.

Figure 1B(iii) shows fluorescence images of the GBM: HUVEC spheroids embedded in a 7.5 mg/mL fibrin gel containing HDF single cells, at day 3 under hypoxic and normoxic conditions. This tri-culture method was found to trigger robust HUVEC angiogenesis creating lumenised capillaries that sprout radially outwards into the surrounding fibrin gel. As in the case of HUVEC-only spheroids, angiogenic sprouting is higher at 20% O_2_ compared to 1% O_2_. Moreover, the GBM cells behave differently depending on whether they are primary or not. NCH82 GBM cells invade into the gel without interacting strongly with the endothelial cells (this is more evident at longer timepoints as shown in Figure S3). On the other hand, primary GBM cells integrate more into the vasculature behaving similarly to the HUVEC. This phenomenon will be explored further in the following sections.

### 2.4. GBM promotes angiogenesis

In order to investigate the effect of glioblastoma cells on the sprouting of endothelial cells, the full tumoroid model (GBM: HUVEC spheroid surrounded by HDF single cells) or the exact model but with-\ out the GBM cells (HUVEC spheroid with HDF) were compared. Figure 2A depicts the angiogenic sprouting of HUVEC spheroids under various co-culture conditions at 1% and 20% O_2_. For each condition, the explant area, the average vessel length and the junction density are presented in Figure 2B. The results show that in the presence of primary glioblastoma cells, endothelial cells produce vasculature of higher explant area but lower interconnectivity. In addition, an increase in the vessel length was observed under hypoxic conditions only (Figure 2B). From the above, we hypothesize an invasive mode of angiogenesis for primary GBM cells. This suggests that the cells are producing tumor specific growth factors. The NCH82 GBM cell line interacts differently with the endothelial cells inducing sprouting within a smaller explant area while the average vessel length remains unaffected. As with the primary GBM, capillaries of lower interconnectivity are produced (Figure 2B).

**Figure 2.**
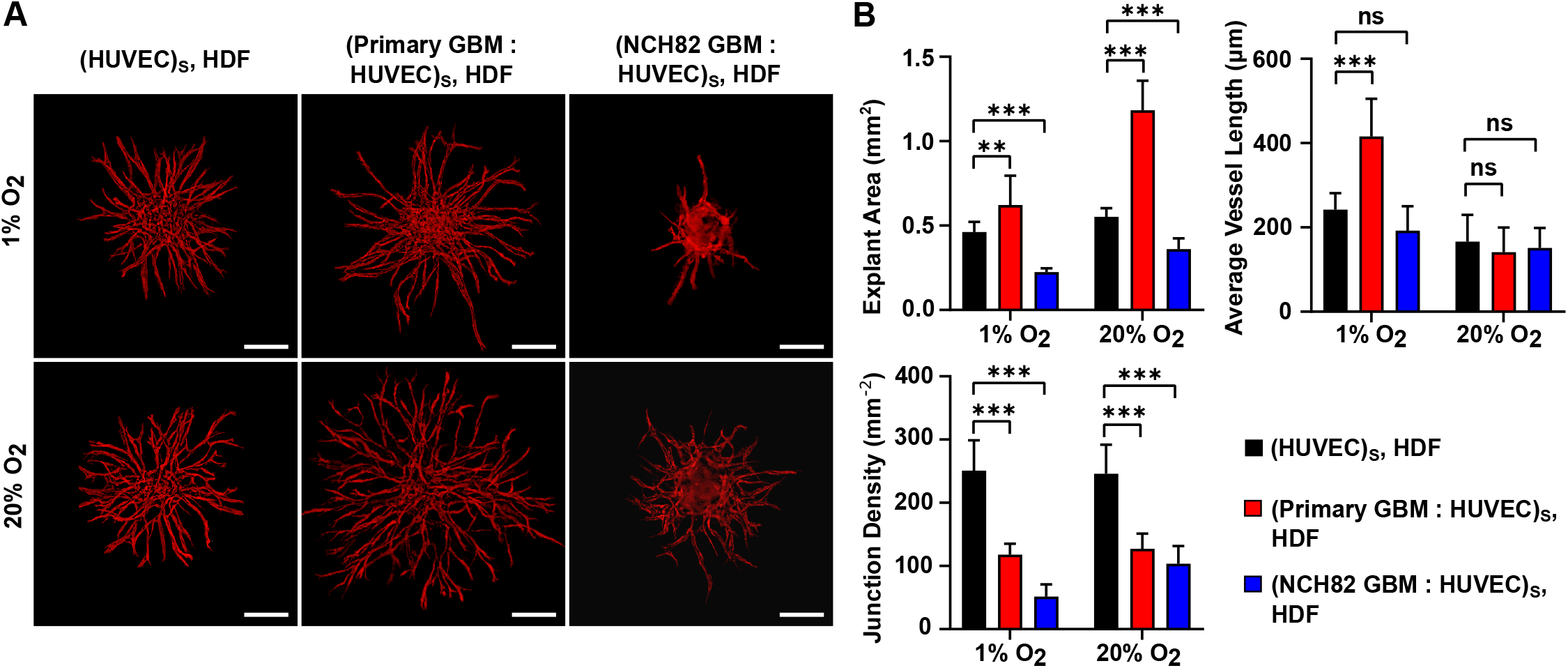
Angiogenic sprouting of HUVEC under various culture conditions at day 3 after seeding cultured at 1% and 20% O_2_. (A) Fluorescence images showing sprouting of HUVEC (red) spheroid with HDF, and primary or NCH82 GBM (green): HUVEC spheroid with HDF. HDF cells are not stained so they are non-fluorescent. The spheroids and single HDF cells are embedded in a 7.5 mg/mL fibrin gel. The scale bars represent 200 *µ*m. (B) Graphical representations of the explant area, average vessel length and junction density. Data are presented as the mean ± standard deviation; *n* = 3; \**p <* 0.05, \*\**p <* 0.01, \*\*\**p <* 0.001. Subscript S denotes cells seeded in the same spheroid.

These results suggest a difference in angiogenic capabilities between primary and NCH82 GBM cells. Patient-derived primary GBM cells have been cultured under conditions that retain their stem-like potential [28, 29] whereas in the serum-grown NCH82 cell line the cells are differentiated. This is consistent with *in vivo* observations as tumors that have a high stem cell number are highly angiogenic [15, 16]. Furthermore, the lower vascular interconnectivity observed in the presence of GBM cells is an important biomimetic feature as it is consistent with the high amount of dead ends present in the GBM vasculature compared to normal vasculature [30, 31].

### 2.5. GBM as supporting cells

In a related experiment we tested the hypothesis that primary GBM cells can act as a supporting cell type for vessel formation, role taken up by the fibroblasts in the previous experiment. For this experiment, a 1000-cell HUVEC spheroid was cultured in a 7.5 mg/mL fibrin gel containing 10^6^ cells/mL primary or NCH82 GBM single cells. The angiogenic sprouting of HUVEC under 1% and 20% O_2_, with or without additional growth factors, is illustrated in Figure 3A for day 3. The number of HUVEC sprouts is shown in Figure 3B.

**Figure 3.**
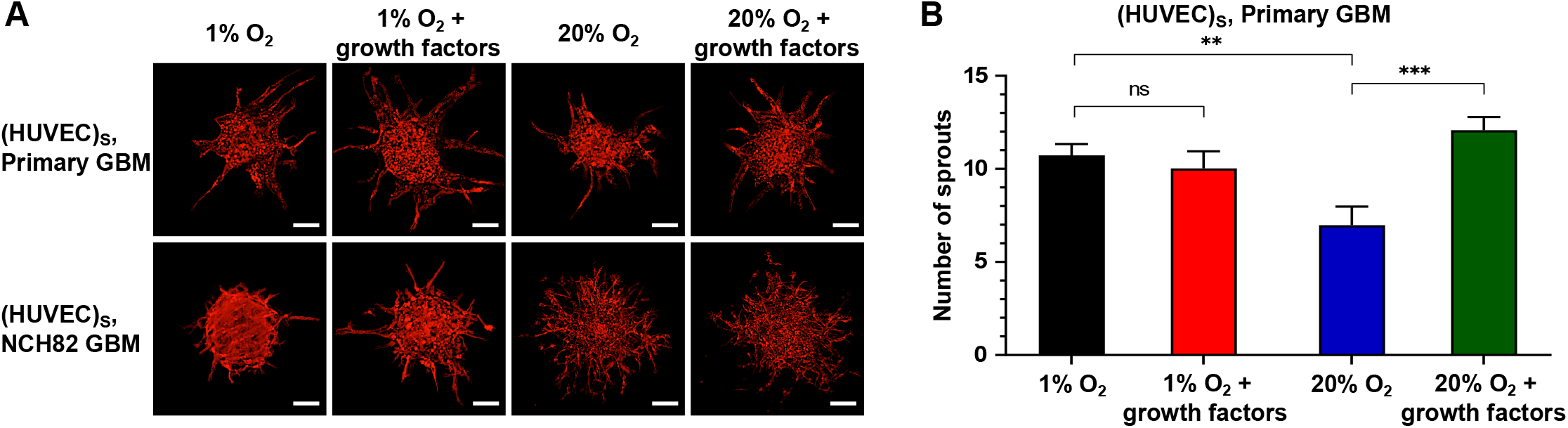
Angiogenic sprouting of HUVEC spheroids with GBM as the supporting cell at 1% and 20% O_2_, with or without additional growth factors. (A) Red fluorescence channel images showing HUVEC spheroids with supporting primary or NCH82 GBM cells. Images were taken at day 5 after seeding. HUVEC are shown in red. GBM single cells are not visible. The spheroids and single GBM cells are embedded in a 7.5 mg/mL fibrin gel. The scale bars represent 100 *µ*m. (B) Graphical representation of the number of HUVEC sprouts. Data are presented as the ± mean standard deviation; *n* = 3; \**p <* 0.05, \*\**p <* 0.01, \*\*\**p <* 0.001. Subscript S denotes cells seeded in the same spheroid.

When primary GBM is used as a supporting cell, endothelial sprouting is more pronounced in hypoxia compared to normoxia (Fig. 3B). The opposite result is observed for HDF supporting cells (Figure 2A), demonstrating the supporting role primary, serum-free cultured GBM cells have especially under hypoxia even without direct intracellular contact. GBM cells are known to secrete pro-angiogenic factors under hypoxia, such as VEGF and bFGF [32, 33].

Futhermore, the addition of exogenous growth factors enhances this effect for cells cultured at normoxia but not at hypoxia suggesting that under hypoxia the GBM cells are already producing a significant amount of growth factors (Figure 3B). In the absence of GBM cells, HUVEC undergo a migratory response which suggests that GBM cells are indeed producing pro-angiogenic factors and these factors recruit cells that may contribute to vessel formation. Performing the same experiment using the NCH82 GBM cells, endothelial cells are found to mostly invade the gel rather than sprout, particularly under normoxia, showing that these cells are not so capable of inducing HUVEC vascularization (Figure 3A).

### 2.6. Endothelial-like behaviour of GBM cells

Time-lapse images of the GBM vascularized tumoroid model depict the evolution of GBM: HUVEC spheroids with HDF single cells at 1% O_2_ over 32 h starting at day 1 after seeding. In a feature unique to primary GBM vascularized tumoroids, tumor cells grow together with the endothelial cells forming sprouting networks. Moreover, as time-lapse imaging shows, some of the formed sprouts are composed of primary GBM cells (Figure 4A). However, in NCH82 vascularised tumoroids, GBM cells break away from the primary mass and invade separately into the surrounding matrix (Figure 4B). The co-localization between the tumor and the endothelial cells is quantified using the Pearson’s correlation coefficient (Figure 4C). The coefficient is steadily high *∼* 0.75 for primary GBM cells but it decreases from *∼* 0.6 to *∼* 0.2 for NCH82 GBM cells over 32 h.

**Figure 4.**
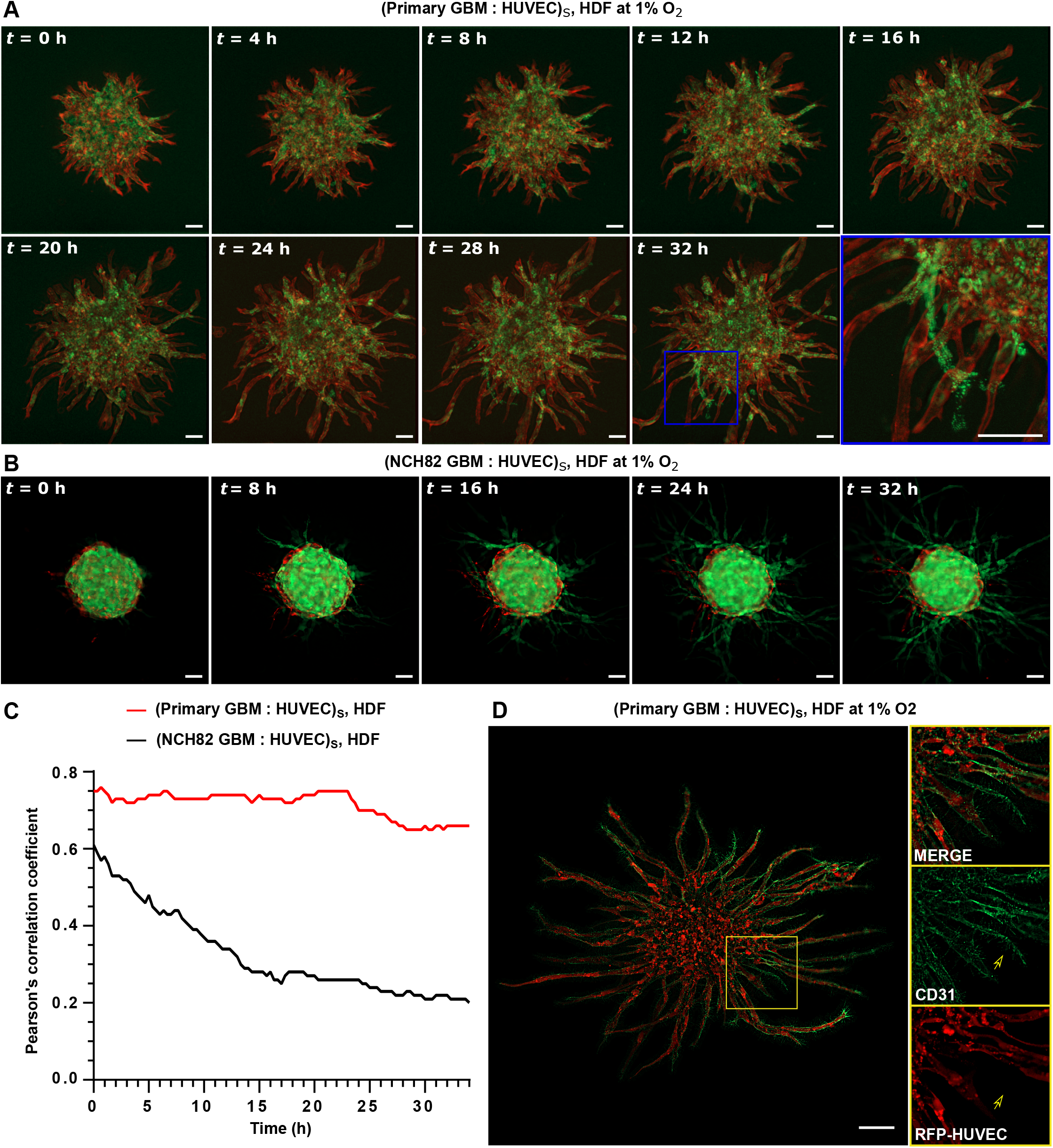
Time-lapse imaging showing evolution of GBM: HUVEC spheroid with HDF as the supporting cell at 1% O_2_ over a period of 32 h. (A) Primary GBM (green): HUVEC (red) spheroid. (B) NCH82 GBM (green): HUVEC (red) spheroid. *t* = 0 h is taken at day 1 after seeding. HDF cells are not stained so they are non-fluorescent. The spheroids and single HDF cells are embedded in a 7.5 mg/mL fibrin gel. (C) Plot showing Pearson’s correlation coefficient as a function of culture time. Similar results were obtained in three independent experiments. (D) CD31 immunohistochemical staining (green) of primary GBM: HUVEC spheroid at day 3 after seeding. Right images are magnified views of the selected area (yellow box). The scale bars represent 100 *µ*m. Subscript S denotes cells seeded in the same spheroid.

CD31 immunohistochemical staining of primary GBM: HUVEC vascularised tumoroids (Figure 4D) revealed a mosaic pattern of the sprouts, showing cells other than HUVEC expressing CD31, suggesting that the primary GBM cells have acquired an endothelial-like behaviour (Figure 4D, arrow in the magnified views of the selected area (yellow box)). This effect was found to be more prominent under hypoxia which is consistent with what has been observed *in vivo* [34, 35].

## 3. Conclusion

A human, disease-relevant, *in vitro* vascularized tumoroid model has been developed to investigate microenvironment cues of glioblastoma (GBM) angiogenesis. The model comprises of GBM and endothelial cell (HUVEC) spheroids in a fibrin gel containing human dermal fibroblasts. It is easily scaled up, tractable, cost-effective, patient-specific, can employ naturally-derived and synthetic matrices, and is amenable to imaging and detailed analysis such as spheroid growth tracking. Although the model lacks perfusable vasculature and has no pre-existing capillary bed or blood brain barrier, it is complex enough to recapitulate features of GBM angiogenesis, which have previously been observed *in vivo*. Patient-derived primary GBM cells were found to support angiogenic sprouting when cultured under conditions that preserve their stem-like potential, in contrast to a GBM cell line containing differentiated cells. Angiogenic sprouting was observed even in the absence of supportive fibroblasts. In the presence of exogenous growth factors, sprouting was enhanced under normoxia but not at hypoxia suggesting that hypoxic GBM cells are already producing a significant amount of growth factors. Primary GBM cells cocultured with HUVEC in spheroids, produced vasculature of higher explant area and lower connectivity in comparison to HUVEC-only spheroids. Under hypoxia, primary GBM cells co-localize with HUVEC to form sprouting vascular networks and even acquire endothelial-like characteristics. These findings demonstrate that our 3D tumoroid *in vitro* model exhibits biomimetic attributes. It therefore holds potential as a simple reductionist model for the development of anti-angiogenic treatments to GBM especially in view of the need to assess the preliminary effectiveness of new treatments before moving to more complex and lengthy efficacy *in vivo* models.

## 4. Experimental Section

### Cell culture

Primary GBM cells from fresh surgical samples (Addenbrooke’s Hospital, UK) were derived using the Cambridge Protocol [29]. Cells were cultured in serum-free medium (SFM) based on phenol red-free Neurobasal A medium (Invitrogen, UK) supplemented with 2 mM L-glutamine and 1% Antibiotic -Antimycotic (Life Tech, UK) with 20 ng/mL human epidermal growth factor (hEGF, Sigma-Aldrich, UK), 20 ng/mL human basic fibroblast growth factor (hbFGF, R & D systems, UK), 2% (v/v) B27 (Invitrogen, UK) and 1% N2 (Invitrogen, UK). They were propagated as monolayers on extracellular matrix-coated flasks (ECM 1:50 dilution, Sigma-Aldrich, UK). Culturing GBM cells under serum-free conditions with hEGF and hbFGF has been reported [36, 37] to tentatively preserve the characteristics of the primary brain tumors.

GBM cell line NCH82, a gift from Prof Conceição Pedroso de Lima (University of Coimbra, Portugal), was transduced with GFP through lentiviral infection. The NCH82 cell line was cultured in serum-containing, high glucose Dulbecco’s Modified Eagle’s Medium (DMEM -high glucose, Life Tech, UK), containing 10% fetal bovine serum (FBS, Life Tech, UK) and 1% antibiotic antimycotic solution (Sigma-Aldrich, UK). Primary and NCH82 GBM cells were passaged at 80% confluency with accutase (Sigma-Aldrich, UK).

Red fluorescent protein expressing HUVEC (2B Scientific,UK) were cultured in Endothelial Growth Medium 2 (EGM2, Promocell, Germany) and 1% antibiotic antimycotic solution. HDF (Public Health England) were cultured in high glucose DMEM containing 10% FBS (Life Technologies, UK) and 1% antibiotic antimycotic solution. HUVEC and HDF were passaged at 80% confluency with TrypLE Express (Life Technologies, UK). All cells were incubated at 37^*°*^C in 5% CO_2_ and 95% relative humidity.

### Multi-cellular spheroid formation

Co-culture spheroids were prepared using the hanging drop method. A stock solution of methylcellulose was prepared by dissolving 1.2 g methycellulose powder (Sigma-Aldrich, UK) in 100 mL of DMEM. GBM and/or HUVEC cells were detached and suspended in 1:1 EGM to SFM medium containing 20% methycellulose (v/v). 20 *µ*L cell suspension drops each containing 1000 cells were pipetted onto a non-adherent petri dish (Greiner, UK). Co-culture spheroids were composed of HUVEC to GBM at a 3:1 ratio. HUVEC-only and GBM-only spheroids were also generated. The dish was turned upside down and incubated at 37^*°*^ C. The spheroids were harvested after 16 h.

### Gel construct preparation

For each fibrin construct, approximately 3 spheroids were embedded in 25 *µ*L of fibrin gel. Fibrin gels contained HDF or GBM at 10^6^ cells/mL or no cells. To prepare the fibrin gel, fibrinogen was dissolved in 1:1 EGM2 to SFM medium, with or without additional pro-angiogenic growth factors (200 ng/mL VEGF and bFGF), to a final concentration of 7.5 mg/mL and was mixed with 1 *µ*L of 50 U/mL thrombin. The solution was pipetted into a 48-well plate, turned upside down, left at room temperature undisturbed for 10 min and then moved to a 37^*°*^C incubator for 20 min to completely polymerise. 1:1 EGM to SFM medium was then added. Fibrin constructs were maintained at either 20% O_2_ (normoxia) or 1% O_2_ (hypoxia). Medium was changed every 2 days.

### Fluorescent labelling

Primary GBM cells were labelled using Qtracker 525 (ThermoFisher Scientific, UK). Cells were subcultured at a density of 2 *×* 10^4^ cells per well of a 48-well plate and incubated overnight. Then 200 *µ*L of 15 nM Qtracker diluted in SFM was added per well. Cells were then incubated for 6 h before washing them twice with SFM.

Tumoroids were immunofluorescently stained with CD31 antibody. Gel constructs were fixed using 4% paraformaldehyde (PFA, Sigma-Aldrich, UK) for 1 hour followed by washes with phosphate-buffered saline (PBS, Sigma-Aldrich, UK). The constructs were subsequently permeabilised with 0.25% Triton X-100 (Sigma-Aldrich, UK) and blocked with 1% bovine serum albumin (BSA, Sigma-Aldrich, UK). Cells were incubated overnight with CD31 antibody (Abcam, UK) diluted at 1:100 followed by 2 h of incubation with 1:500 AlexaFluor 568 goat anti-mouse IgG (abcam, UK) diluted at 1:500. Washes with PBS followed both incubation periods.

### Visualization and quantification of sprouting angiogenesis

Images of sprouting tumoroids were taken using epi-fluorescence microscopy (Zeiss Axio Observer Z1 inverted microscope with an ORCA-Flash4.0 camera). The microscope was equipped with an incubator regulating CO_2_ (5%), O_2_ (20% or 1%), temperature (37^*°*^C) and humidity (95%) (Okolab, Italy). Time-lapse images were acquired every 20 min. The images were the result of the deconvolution of an image z-stack using ZEN software (Zeiss, Germany).

The area of the vessel networks that extended beyond the surface of the initial spheroid boundary was selected using ImageJ. This area was then analysed using AngioTool [38]. The software provided automatic measurements of: (i) the total vessel area, (ii) the total vessel length, (iii) the explant area (area of convex hull containing all the vessels), (iv) the average vessel length and (v) the junction density (total number of vessel junctions normalised by the area). A vessel was defined as the segment between two junction points or a junction point and an end point. The total spheroid area (area of spheroid including any sprouting) was quantified in ImageJ.

GBM co-localization along blood vessels was quantified by the Pearson’s correlation coefficient in Coloc 2 [39] (ImageJ plugin [40]).

### Statistical analysis

Data comparisons between two sets of data were performed in GraphPad Prism 8 software (GraphPad, San Diego, USA). Data are displayed as mean ± standard deviation. Three spheroids were analysed out of each one of three independent experiments. The explant area of each spheroid ranged between 0.1 and 1.3 mm^2^. Two sets of data were compared using the Students t-test. The threshold for statistical significance was set at a value of \**p <* 0.05. \*\**p <* 0.01; \*\*\**p <* 0.001.

## Acknowledgements

This research was supported by an Engineering for Clinical Practice grant (to AEM and CW) and an EPSRC IAA Follow on Fund (EP/R511675/1 to AEM). Financial support for AST and AWJ has been provided via an WD Armstrong studentship and EPSRC (EP/R511675/1), respectively. The authors would like to thank Dr Rasha Rezk (University of Cambridge, UK) and Prof Conceiçaõ Pedroso de Lima (University of Coimbra, Portugal) for providing the patient-derived primary GBM cells and the NCH82 GBM cell line, respectively. We also thank Dr John Ong (University of Cambridge, UK) for helpful discussions on antibodies.

## Data availability

The datasets generated and analysed during the current study are available to download from Mendeley Data, V1, doi: 10.17632/kc64v677tb.1.

