## Supplementary Materials for "A Vascularized Tumoroid Model for Human Glioblastoma Angiogenesis" (from another version of this preprint: version 1, 2021-02-07)

### Supplementary material

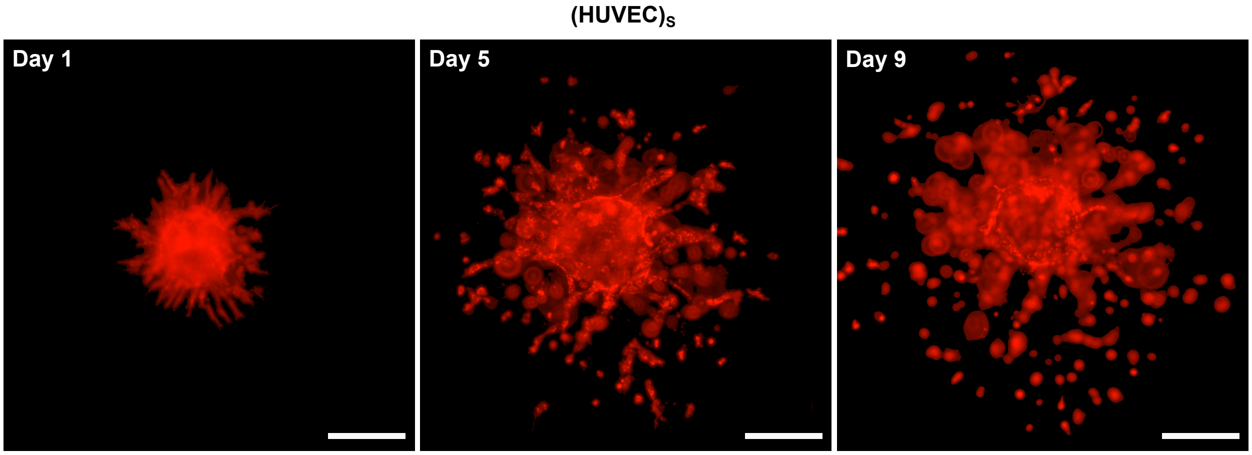


**Figure 1.** Fluorescence images showing the evolution over time (days 1, 5 and 9) of a HUVEC spheroid embedded in a 7.5 mg/mL fibrin gel at 20% $O_2$. The scale bars represent 250 $\mu$m. Subscript S denotes cells seeded in the same spheroid.

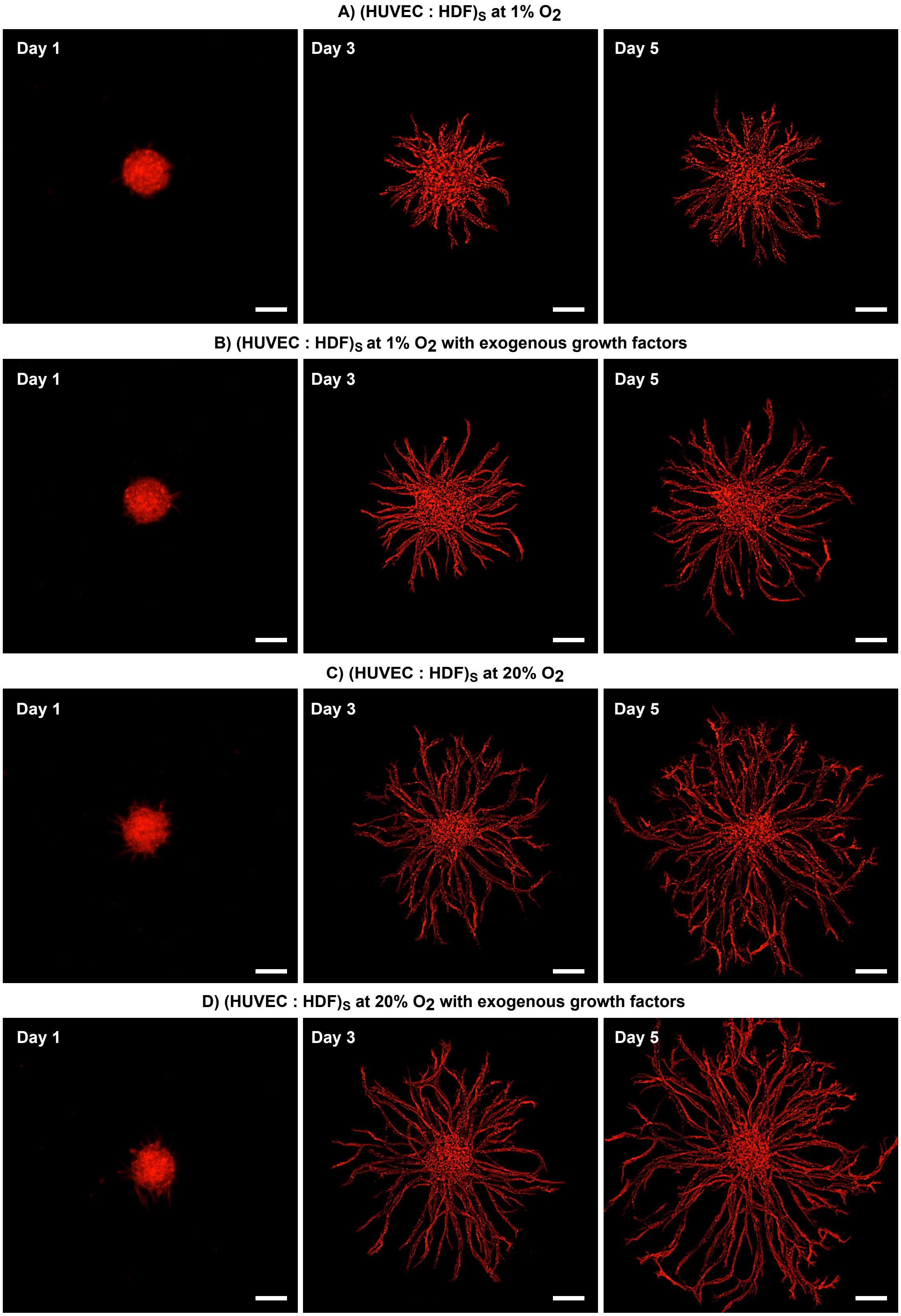


**Figure 2.** Fluorescence images showing the evolution over time (days 1, 3 and 5) of a HUVEC (red) spheroid surrounded by single HDF cells in a 7.5 mg/mL fibrin gel at (i) 1% $O_2$, (ii) 1% $O_2$ with exogenous growth factors, (iii) 20% $O_2$ and (iv) 20% $O_2$ with exogenous growth factors. HDF cells are not stained so they are non-fluorescent. The scale bars represent 250 $\mu$m. Subscript S denotes cells seeded in the same spheroid.

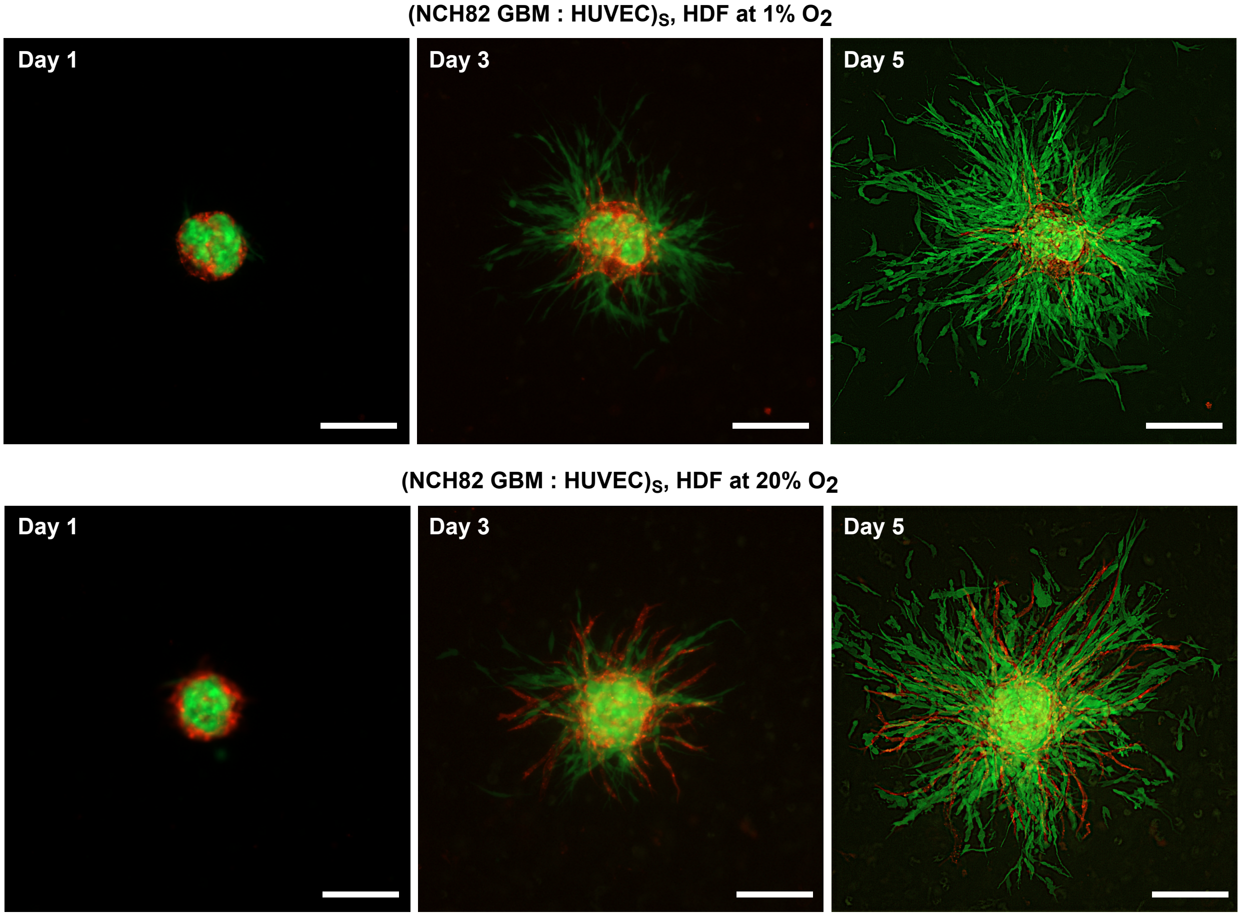


**Figure 3.** Fluorescence images showing the evolution over time (days 1, 3 and 5) of a NCH82 GBM (green) : HUVEC (red) spheroid surrounded by single HDF cells in a 7.5 mg/mL fibrin gel at 1% and 20% $O_2$. HDF cells are not stained so they are non-fluorescent. The scale bars represent 250 $\mu$m. Subscript S denotes cells seeded in the same spheroid.
